# Engineering a Vaccine Platform using Rotavirus A to Express SARS-CoV-2 Spike Epitopes

**DOI:** 10.1101/2022.03.23.485570

**Authors:** Ola Diebold, Victoria Gonzalez, Luca Venditti, Colin Sharp, Rosemary A. Blake, Joanne Stevens, Sarah Caddy, Paul Digard, Alexander Borodavka, Eleanor Gaunt

**Affiliations:** Infection and Immunity Division, Roslin Institute, University of Edinburgh, Easter Bush Campus, Midlothian, EH25 9RG, United Kingdom; Department of Biochemistry, University of Cambridge, UK; Cambridge Institute of Therapeutic Immunology and Infectious Disease, Department of Medicine, University of Cambridge, UK

**Keywords:** rotavirus, NSP3, VP4, reverse genetics

## Abstract

Human rotavirus (RV) vaccines used worldwide have been developed using live attenuated platforms. The recent development of a reverse genetics system for RVs has delivered the possibility of engineering chimeric viruses expressing heterologous peptides from other virus species to generate polyvalent vaccines. We tested the feasibility of this using two approaches. Firstly, we inserted short SARS-CoV-2 spike peptides into the hypervariable region of the simian SA11 RV strain viral protein (VP) 4. Secondly, we fused the receptor binding domain (RBD) of the SARS-CoV-2 spike protein, or the shorter receptor binding motif (RBM) nested within the RBD, to the C-terminus of non-structural protein (NSP) 3 of the bovine RF strain RV, with or without an intervening T2A peptide. Mutating the hypervariable region of SA11 VP4 impeded viral replication, and for these mutants no cross-reactivity with spike antibodies was detected. To rescue NSP3 mutants, we established a plasmid-based reverse genetics system for the bovine RF strain. Except for the RBD mutant, all NSP3 mutants delivered endpoint titres and replication kinetics comparable to that of the WT virus. In ELISAs, cell lysates of an NSP3 mutant expressing the RBD peptide showed cross reactivity with a SARS-CoV-2 RBD antibody. 3D bovine gut enteroids were susceptible to infection by all NSP3 mutants but only RBM mutant showed cross reactivity with SARS-CoV-2 RBD antibody. The tolerability of large peptide insertions in the NSP3 segment highlights the potential for this approach in the development of vaccine vectors targeting multiple enteric pathogens simultaneously.

**IMPORTANCE:** We explored the use of rotaviruses (RVs) to express heterologous peptides, using SARS-CoV-2 as an exemplar. Small SARS-CoV-2 peptide insertion (<34 amino acids) into the hypervariable region of the viral protein 4 (VP4) of RV SA11 strain resulted in reduced viral titre and replication, thus limiting its use as a potential vaccine expression platform. To test RF strain for its tolerance for peptide insertions, we constructed a reverse genetics system. NSP3 was C-terminally tagged with SARS-CoV-2 spike peptides of up to 193 amino acids. With a T2A-separated 193 amino acid tag on NSP3, there was little effect on the viral rescue efficiency, titre and replication. Tagged NSP3 elicited cross-reactivity with SARS-CoV-2 spike antibodies in ELISA. This is the first report describing epitope tagging of VP4, and of a reverse genetics system for the RF strain. We highlight the potential for development of RV vaccine vectors targeting multiple enteric pathogens simultaneously.

## INTRODUCTION

Group A rotaviruses (RVs) are a leading cause of severe acute gastroenteritis in infants and young children worldwide, accounting for ~215,000 deaths annually [1–3]. Likewise, RV-associated enteritis in young calves and piglets has a significant economic impact on livestock production as a result of the high morbidity and mortality caused [4–7]. Two human live attenuated RV vaccines, Rotarix and RotaTeq have proven effective in reducing the incidence of RV-related hospitalisation and mortality internationally [8–11]. Vaccination strategies for livestock rely on induction of active or passive immunity using animal RV vaccines [12–15].

RV is a double-stranded RNA (dsRNA) virus with 11 genome segments encoding six structural viral proteins (VP1 - VP4, VP6 and VP7) and depending on the strain, five or six non-structural proteins (NSP1 - NSP5 ± NSP6) that are essential for virus replication [1, 16–18]. The mature infectious virion, termed a triple-layered particle (TLP), consists of an outer layer formed by VP4 and VP7. A double-layered particle (DLP), nested within the TLP, contains the intermediate and inner layers of the capsid [1]. RV primarily infects mature enterocytes of the intestinal epithelium and replicates exclusively in the cytoplasm [19, 20]. Efficient RV cell entry requires proteolytic cleavage of the outer capsid protein, VP4, into VP8* (28kDa) and VP5* (60kDa) domains by trypsin-like proteases of the host gastrointestinal tract [21–23]. The VP8* lectin domain mediates RV attachment to different host cell receptors such as sialic acid-containing glycans, histo-blood group antigens and integrins, depending on the virus strain [20, 24–26]. Low calcium levels trigger dissociation of VP7 and VP4, releasing the transcriptionally active DLP into the cytoplasm [1]. Here, DLPs transcribe capped, non-polyadenylated, positive-sense single-stranded RNA transcripts, which act as templates for viral protein translation [1]. The 11 mRNAs share a conserved terminal 3’-GACC sequence that contains *cis*-acting signals important for transcription by the RNA-dependent RNA polymerase (VP1) [27–31]. The 3’ GACC sequence is bound by the N-terminus of NSP3, while the C-terminus of NSP3 interacts with the eukaryotic protein initiation factor 4GI (eIF4GI) [32–34]. NSP3 displaces poly-A binding protein from ribosomal complexes to favour the translation of viral over host cell proteins [35–37].

The molecular characterisation of RVs has historically proven challenging due to inefficiencies of the helper virus-dependent reverse genetics systems available for controlled mutagenesis studies [38–41]. Recently however, Kanai *et al*. developed a plasmid-only reverse genetics system for the simian RV strain SA11 [42]. The plasmid-only reverse genetics system has been optimised to study the functions of RV proteins, to generate RV reassortants and RV reporter expression systems, and conceptualise novel vaccine platforms [43–45]. Using the plasmid-only reverse genetics system, recombinant RVs harbouring chimeric VP4 proteins that showed efficient replication in cell culture and neutralising activity in mice have also been engineered [46–48]. Protection from infection with RV is primarily mediated by heterotypic neutralising antibodies that target VP4 and/ or VP7. VP4 is therefore highly immunogenic and an important target for adaptive immunity [49, 50]. Thus, the potential for VP4 to express heterologous epitopes from different RV strains may provide a delivery platform for expression of different vaccine antigens, though peptide insertions into VP4 has not previously been tested. Additionally, the plasmid-only reverse genetics system has been utilised to generate a repertoire of recombinant RVs expressing fluorescent reporter proteins [48, 51]. The C-terminus of the SA11 NSP3 open reading frame (ORF) was fused to a porcine teschovirus translational 2A element followed by various reporters including UnaG, mKate, mRuby or TagBFP to successfully yield two uncoupled proteins without compromising virus replication [51]. A more recent study showed that the C-terminus of SA11 NSP3 can express different peptides of the severe acute respiratory syndrome coronavirus 2 (SARS-CoV-2) spike protein with minimal impact on endpoint titres [52]. This has highlighted the potential to use RVs as expression vectors for development of polyvalent vaccines for enteric viruses.

In response to the recent emergence of SARS-CoV-2, the causative agent of coronavirus disease 2019 (COVID-19), several platforms have been utilised for vast global vaccine production [53, 54]. Current licensed vaccines include examples that employ mRNA (Pfizer-BioNTech, Moderna) or viral vectors (AstraZeneca, Janssen) to deliver genetic material encoding the SARS-CoV-2 spike protein in order to stimulate the production of neutralising antibodies and T-cell mediated immune responses that target this protein [55–58]. High neutralising antibody titres are strongly associated with the receptor binding domain (RBD) of spike, making it the most immunogenic antigen [59–62]. To assess the potential for generating chimeric vaccines using RV, we used SARS-CoV-2 as a timely exemplar to introduce spike peptides into an RV backbone and determine whether chimeric viruses showed cross-reactivity with spike antibodies. For this, the hypervariable region of SA11 VP4 (VP8* lectin domain) and the C-terminus of the bovine RF strain NSP3 were modified to express SARS-CoV-2 spike epitopes.

This is the first report describing tagging of the surface protein VP4 of the RV with heterologous peptides. We found that mutating the hypervariable region of SA11 VP4 reduced RV infectivity and mutants expressing spike peptides did not cross-react with SARS-CoV-2 spike antibodies, suggesting that VP4 tagging is not a viable strategy for live attenuated vaccine development. Using our established reverse genetics system for the bovine RF strain RV, we rescued infectious viruses expressing either the RBD or the RBM of the SARS-CoV-2 spike protein, with similar titres and replication kinetics to those of the wild type (WT) virus. These viruses cross-reacted with RBD antibodies in ELISA, and were able to infect bovine gut enteroids, inferring the potential of the system for use in live attenuated vaccine development.

## MATERIALS AND METHODS

### Cell lines

African Green monkey kidney epithelial (MA104) cells were cultured in Dulbecco’s modified Eagle’s medium (DMEM) (Sigma-Aldrich) supplemented with heat inactivated 10% foetal bovine serum (FBS) (Gibco) and 1% penicillin-streptomycin (Gibco). Cells of the BSR-T7 clone of baby hamster kidney fibroblasts (BHK-21 cells), constitutively expressing T7 RNA polymerase, were cultured in Glasgow’s Minimal Essential Medium (GMEM) (Gibco) supplemented with 1% tryptose phosphate broth (TPB) (Gibco), heat inactivated 10% FBS (Gibco) and 1% penicillin-streptomycin (Gibco). Both cell lines were a kind gift from the laboratory of Prof. Massimo Palmarini (MRC-University of Glasgow Centre for Virus Research, UK). Cells were passaged twice weekly and maintained at 37°C, 5% CO_2_.

### Design of RV expression vectors containing spike epitopes

To engineer mutant RVs expressing SARS-CoV-2 spike peptides, we utilised two strategies involving both the simian RV strain SA11 and the bovine RV strain RF. It was previously suggested that the surface protein VP4 of the SA11 strain tolerates short immunogenic peptides inserted at specific sites with minimal impact on viral replication or particle assembly [63]. Guided by the available structure model of VP4 (PDB: 4V7Q), we have chosen an exposed loop outside the sialic acid-binding domain located within the ‘head’ of the VP4 spike (Fig. 1A) to investigate the possibility of VP4 alteration. Moreover, several neutralisation escape mutants were ascribed to the amino acid changes within this region [64], consistent with its accessibility to antibodies. A number of B-cell linear epitopes derived from either the heptad repeat 2 (HR2), N-terminal domain (NTD) or RBM regions of the SARS-CoV-2 spike protein were selected for insertion [61, 65, 66] (Fig. 1A). These epitopes were introduced into the hypervariable region of SA11 VP4 between amino acid position 164 and 198 with linker sequences [67, 68] to increase their accessibility. VP4 plasmid constructs expressing SARS-CoV-2 spike peptides were generated by PCR and denoted HR2, NTD, RBM.1 and RBM.2. These epitopes have been curated from the scientific literature by the Immune Epitope Database available at VIPR (https://www.viprbrc.org/brc/). Only unstructured epitopes of up to 15 residues were chosen (Table 1). The SA11 strain was selected for this mutagenesis due to the structural characterisation of VP4 of its close relative RRV, its user-friendly reverse genetics system and its rapid growth kinetics.

**Figure 1.**
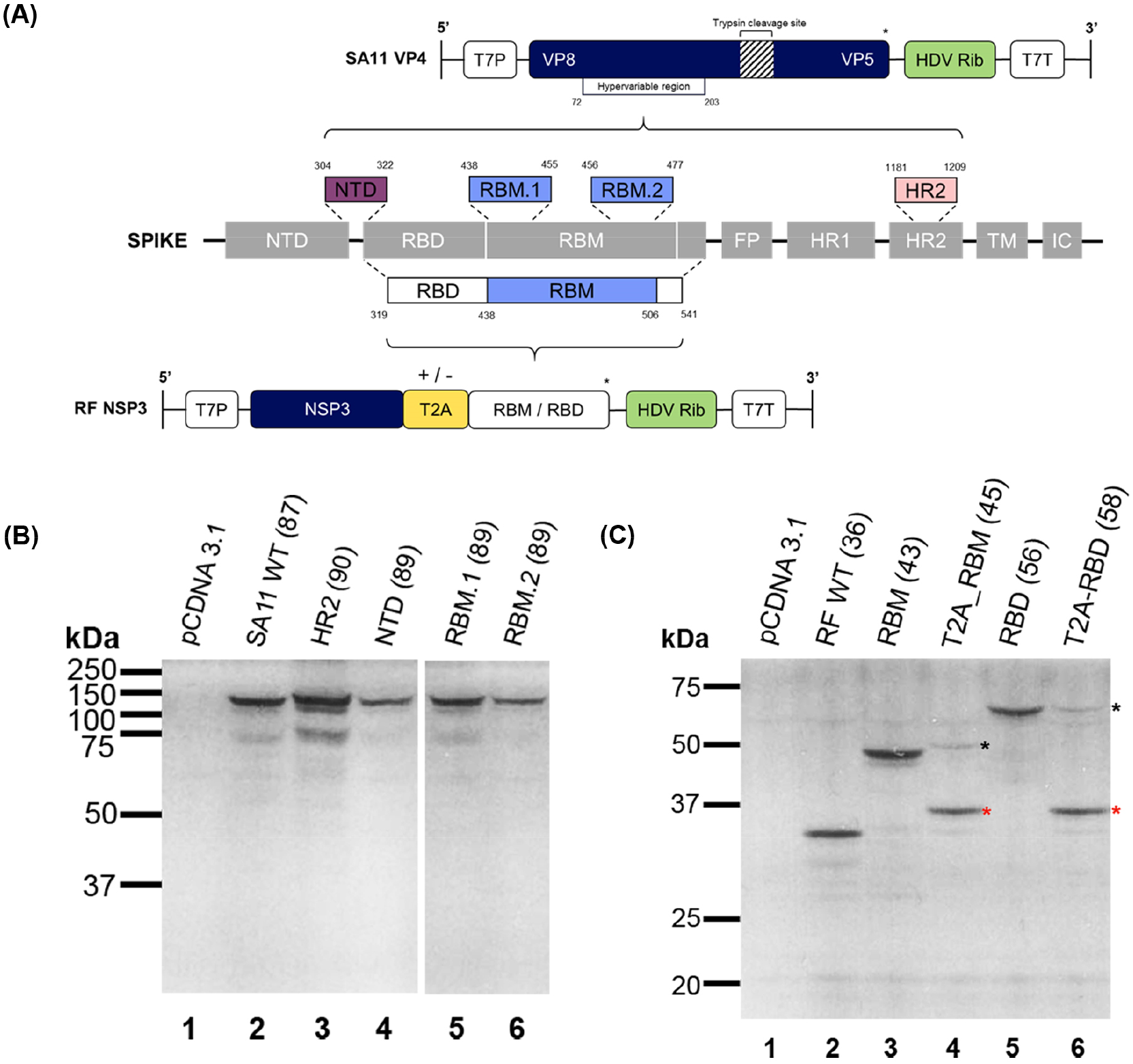
Design and validation of RV VP4 and NSP3 plasmid constructs used to generate mutant viruses. **(A)** Schematic showing overall topology of SARS-CoV-2 spike protein in grey boxes: N-terminal domain (NTD), receptor binding domain (RBD) which contains the receptor binding motif (RBM), fusion peptide (FP), heptad repeats 1 and 2 (HR1 and HR2), transmembrane region (TM) and the intracellular domain (IC) *(Adapted from Lan et al., 2020)* [62]. Dashed lines represent selected regions of the spike protein (coloured boxes with their relative nucleotide positions) that were inserted into the hypervariable region of the SA11 VP4 gene (panel above) and the C-terminus of the RF NSP3 gene (panel below). SA11 VP4 was edited to incorporate either NTD, RBM.1, RBM.2 or HR2 spike peptide sequences. RF NSP3 was fused with either the RBD or RBM sequence with or without T2A (yellow box), represented by +/- sign. Both gene segments were flanked by the T7 promoter (T7P) and the antigenomic HDV ribozyme (‘HDV Rib’, green boxes) followed by T7 terminator (T7T). Asterisks (*) represent stop codons. Schematic not to scale. Coupled *in vitro* transcription and translation reactions of mutated SA11 VP4 **(B)** and RF NSP3 **(C)** segments were carried out using the TnT rabbit reticulocyte lysate system supplemented with [^35^S] methionine. Samples were analysed using SDS-PAGE and autoradiography. The molecular weight marker and the expected product sizes of each segment (in brackets) are indicated (kDa). Empty pCDNA 3.1 vector was used as a negative control. In **(C)**, black asterisks indicate T2A read-through product and red asterisks identify separated products.

**Table 1.**
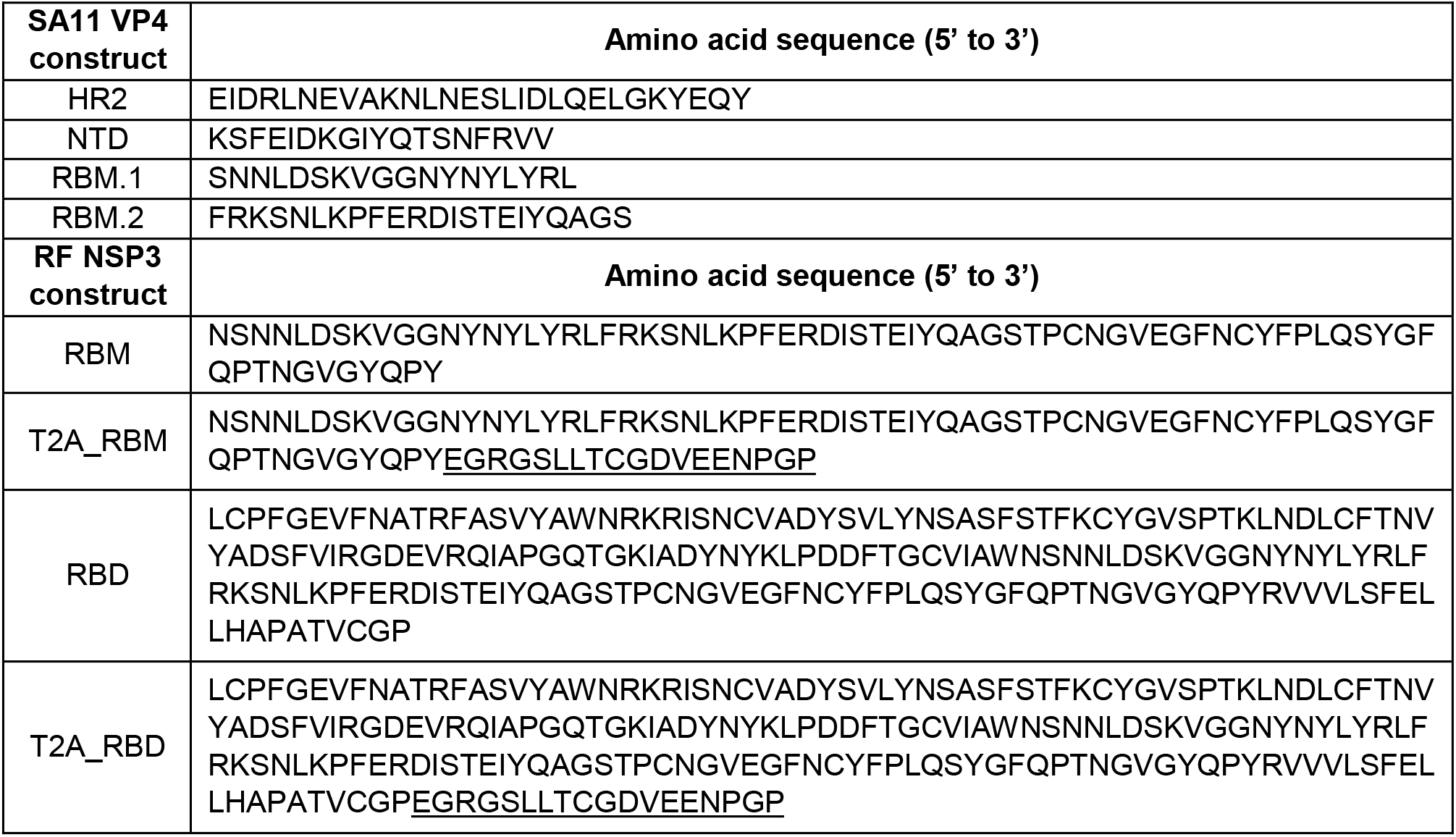
SARS-CoV-2 spike amino acid sequences selected for insertion into the hypervariable region of SA11 VP4 and tagging of RF NSP3. Sequence of the T2A peptide is <u>underlined</u>.

As an alternative to VP4 tagging, we aimed to test whether tagging a RV strain more closely related to the bovine virus backbone used in the pentavalent RotaTeq vaccine [69], could be extrapolated to the bovine RF strain. Here we modified the C-terminus of the RF strain NSP3 ORF to express spike epitopes coding for the RBD or RBM of SARS-CoV-2, with or without an intervening thosea asigna virus 2A (T2A) peptide (Fig. 1A). The inclusion of the T2A was employed in order to lower the risk of interfering with the function of the NSP3 gene. On the other hand, increasing the segment size by incorporating the T2A peptide could further affect the antigenic processing, hence both approaches were trialled. The panel of constructs was assigned the notation RBM, T2A-RBM, RBD and T2A-RBD.

### Plasmid construction

pT7 plasmids used for reverse genetics of SA11 RV were kindly provided by Takeshi Kobayashi [42] through the Addgene plasmid repository against IDs #89162-72. To generate plasmids used for reverse genetics of the bovine RV strain RF, constructs were designed to encode each of the 11 RF gene segments, flanked at the 5’ end by a T7 promoter (T7P) and at the 3’ end an antigenomic hepatitis delta virus (HDV) ribozyme sequences, followed by the T7 terminator sequence (T7T) as in Kanai *et al*. [42]. The constructs were synthesised by Invitrogen GeneArt on either pMK-RQ (kanamycin resistance), pMA-RQ or pMA-T (ampicillin resistance) vectors. RF strain NSP3 constructs RBM, T2A-RBM, RBD and T2A-RBD were ordered as gene blocks from Invitrogen GeneArt and cloned into pT7-NSP2SA11 expression plasmid (Addgene #89169) after the NSP2 ORF was removed using SmaI and SalI restriction enzymes. All plasmids were amplified by transformation into chemically competent *E. coli* DH5α, except RF VP7 encoding plasmid for which DH10β cells were used, and purified using QIAGEN^®^ Plasmid Midi Kit (QIAGEN) according to the manufacturer’s protocol. The presence and the size of the mutation in each plasmid was verified by Sanger sequencing (GATC Biotech or Genewiz, Germany) using primers listed in Table 2. Sequence results were analysed in SSE v1.2 software [70].

**Table 2.**
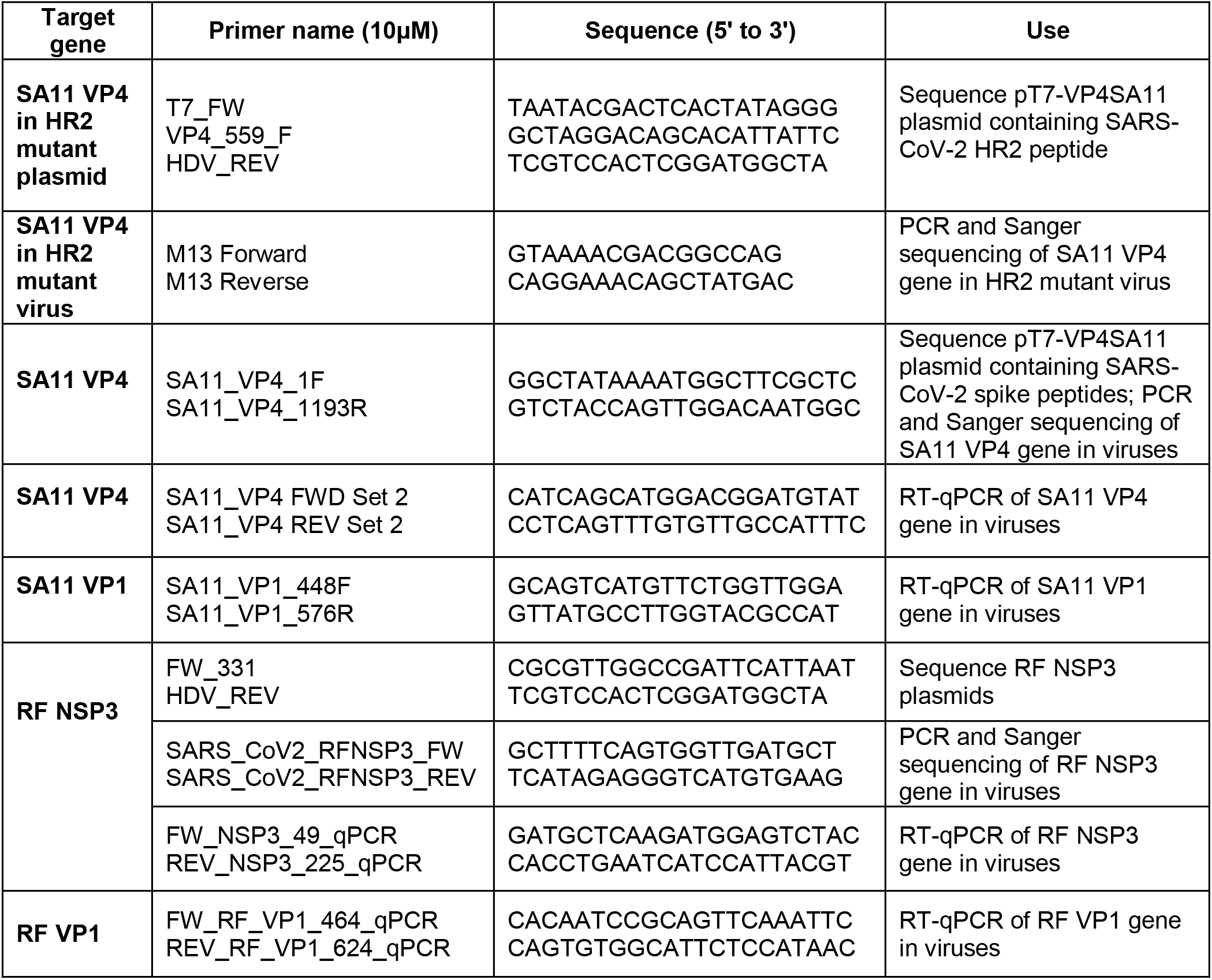
Names and sequences of primers used in this study.

### *In vitro* transcription and translation assay

Coupled *in vitro* transcription and translation reactions were carried out using the Promega TnT® Coupled Reticulate Lysate System labelled with radioactive [^35^S] methionine (PerkinElmer Inc.) according to the manufacturer’s protocol. Briefly, TnT reactions were set up as follows: 8μL TnT mix, 1μCi [^35^S] methionine, 200ng plasmid DNA, made up to 10μL with H_2_O. Mixes were prepared on ice before reactions were incubated at 30°C for 90 min. The reactions were denatured in 8.5μL of 2X Laemmli buffer (65.8mM Tris-HCl [pH 6.8], 100mM DTT [pH 6.8], 2.1% sodium dodecyl sulphate (SDS), 26.3% (w/v) glycerol, 0.01% bromophenol blue) and boiled for 10 min at 95°C. Samples were analysed using sodium dodecyl sulphate–polyacrylamide gel electrophoresis (SDS-PAGE) and autoradiography.

### Autoradiography of dried polyacrylamide gels

Gels were fixed in gel fixing solution (50% (v/v) methanol in water with 10% (v/v) acetic acid) on a rocker for 45 min with the gel fixing solution replaced every 15 min. Fixed gels were transferred onto 3MM Whatman filter paper (Scientific Laboratory Supplies), covered with cling film and dried in a gel dryer (Model 543, Bio-Rad) by heating up to 80°C for 2 hr under vacuum. Dried gels were placed in a sealed cassette with an X-ray film (Fisher Scientific) overnight. X-ray films were developed using a Konica SRX-101A X-ograph film processor following manufacturer’s protocol.

### Reverse genetics system

Viruses were recovered using the protocols described by Kanai *et al*. [42] and Komoto *et al*. [43], with slight modifications. At 70% confluency, monolayers of BSR-T7 cells in 6-well plates were co-transfected with 11 plasmids corresponding to each RV genome segment (2.5μg for plasmids encoding NSP2 and NSP5; 0.8μg for the remaining plasmids) and plasmids encoding two vaccinia virus capping enzyme subunits (pCAG-D1R and pCAG-D12L −0.8μg each) using 16μL Lipofectamine 2000 (Invitrogen) per transfection reaction in a total volume of 200μL of Opti-MEM (Gibco). After 24 hr incubation at 37°C 5% CO_2_, MA104 cells (1 x 10^5^ cells/well) were added to transfected BSR-T7 cells and co-cultured for 4 days in FBS-free DMEM supplemented with 0.5μg/mL porcine pancreatic trypsin type IX (Sigma-Aldrich). Co-cultured cells were then lysed three times by freeze/thaw and lysates were incubated with trypsin at a final concentration of 10μg/mL for 30 min at 37°C 5% CO_2_ to activate the virus. Lysates were then transferred to fresh MA104 cells in T25 flasks and incubated at 37°C 5% CO_2_ for 1 hr. After adsorption, MA104 cells were washed and cultured in FBS-free DMEM supplemented with 0.5μg/mL trypsin type IX for up to 7 days or until complete cytopathic effect was observed. Cells were then lysed three times by freeze/thaw, pelleted and virus-containing supernatants (P1 stocks) were aliquoted and stored at −80°C. To generate mutant SA11 or RF mutant viruses, plasmids encoding either the SA11 VP4 or RF NSP3 gene segment were replaced with the corresponding plasmid encoding SARS-CoV-2 spike epitopes. Mock preparations with the mutated segment omitted were generated for use as negative controls throughout. All rescue experiments were performed three times for each virus panel. The panels of viruses were titred by plaque assays, and the presence of mutations in the target gene segments was confirmed by RT-PCR and Sanger sequencing (GATC Biotech or Genewiz, Germany) as described below. Properties of mutant viruses are summarised in Table 3.

**Table 3.**
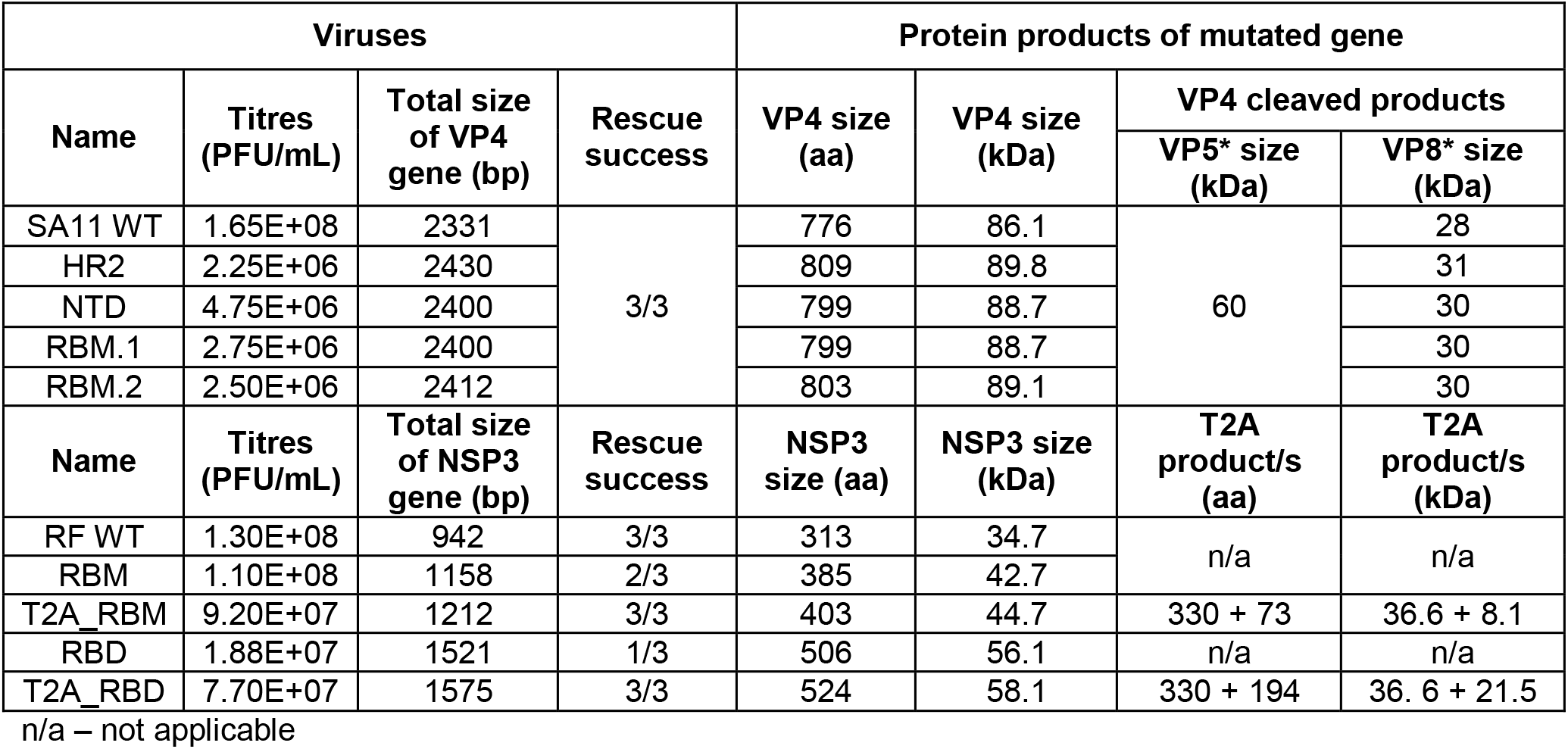
Properties of VP4 and NSP3 mutant viruses. The hypervariable region of SA11 VP4 was tagged with SARS-CoV-2 spike peptides: HR2, NTD and RBM regions. The C-terminus of RF NSP3 was tagged with either the RBM or RBD of SARS-CoV-2 spike with or without a T2A peptide.

| Viruses |  |  |  | Protein products of mutated gene |  |  |  |
| --- | --- | --- | --- | --- | --- | --- | --- |
| Name | Titres (PFU/mL) | Total size of VP4 gene (bp) | Rescue success | VP4 size (aa) | VP4 size (kDa) | VP4 cleaved products |  |
|  |  |  |  |  |  | VP5* size (kDa) | VP8* size (kDa) |
| SA11 WT | 1.65E+08 | 2331 | 3/3 | 776 | 86.1 | 60 | 28 |
| HR2 | 2.25E+06 | 2430 |  | 809 | 89.8 |  | 31 |
| NTD | 4.75E+06 | 2400 |  | 799 | 88.7 |  | 30 |
| RBM.1 | 2.75E+06 | 2400 |  | 799 | 88.7 |  | 30 |
| RBM.2 | 2.50E+06 | 2412 |  | 803 | 89.1 |  | 30 |
| Name | Titres (PFU/mL) | Total size of NSP3 gene (bp) | Rescue success | NSP3 size (aa) | NSP3 size (kDa) | T2A product/s (aa) | T2A product/s (kDa) |
| RF WT | 1.30E+08 | 942 | 3/3 | 313 | 34.7 | n/a | n/a |
| RBM | 1.10E+08 | 1158 | 2/3 | 385 | 42.7 |  |  |
| T2A_RBM | 9.20E+07 | 1212 | 3/3 | 403 | 44.7 | 330 + 73 | 36.6 + 8.1 |
| RBD | 1.88E+07 | 1521 | 1/3 | 506 | 56.1 | n/a | n/a |
| T2A_RBD | 7.70E+07 | 1575 | 3/3 | 524 | 58.1 | 330 + 194 | 36.6 + 21.5 |

n/a – not applicable

### RT-PCR

RNA was purified from viral stocks using the spin protocol of the QIAamp Viral RNA Mini Kit (QIAGEN) followed by RQ1 RNase-Free DNase treatment (Promega) to remove possible plasmid contamination according to the manufacturer’s specifications. cDNA was synthesised with SuperScript™ III Reverse Transcriptase (RT) kit (Invitrogen) using 5μL RNA and 1μL random hexamer primers (10μM) according to the manufacturer’s protocol. PCR was then used to amplify across the regions containing the SARS-CoV-2 spike epitopes in SA11 VP4 or RF NSP3 with primers listed in Table 2 (amplicon product sizes ~1.2-1.4kbp). Using the Q5® High-Fidelity DNA Polymerase PCR Kit (New England BioLabs) for SA11 VP4 and Platinum™ *Taq* DNA polymerase (Invitrogen) for RF NSP3, PCR reactions were set up according to the manufacturer’s instructions in a T100™ Thermal Cycler (Bio-Rad). SA11 VP4 PCR conditions were 1 cycle of 98°C for 30 sec, followed by 35 cycles of 20 sec at 98°C, 20 sec at 52°C and 2 min at 72°C, finishing with a 2 min incubation at 72°C. RF NSP3 PCR conditions were 1 cycle of 5 min at 95°C, followed by 35 cycles of 30 sec at 95°C, 30 sec at 55°C and 2 min at 72°C, finishing with a 5 min incubation at 72°C. PCR product length was confirmed by 0.8% agarose gel electrophoresis containing SYBR™ Safe DNA Gel Stain and gels were imaged using the Odyssey ® XF imaging system (LI-COR). Analyses were performed with Image Studio™ Lite software (LI-COR). The presence of sequence insertions was confirmed by sequencing at GATC Biotech or Genewiz, Germany using primers listed in Table 2. Sequence results were analysed in SSE v1.2 software [70].

### Plaque assay

Plaque assays for RVs were performed using adapted methods [71, 72]. Confluent monolayers of MA104 cells in 6-well plates were washed with FBS-free DMEM and infected with 800μL of ten-fold serially diluted virus for 1 hr at 37°C 5% CO_2_. Following virus adsorption, 2mL/well overlay medium was added (1:1 ratio of 2.4% Avicel (FMC Biopolymer) and FBS-free DMEM supplemented with 0.5μg/mL trypsin type IX) and incubated for 4 days. Cells were then fixed for 1 hr with 1mL/well of 10% neutral buffered formalin (NBF) (CellPath) and stained for 1 hr with 0.1% Toluidine blue (Sigma-Aldrich) dissolved in H_2_O.

### Multi-step virus growth kinetics

To compare growth kinetics of WT and mutant viruses, monolayers of MA104 cells at 70% confluency in 24-well plates were infected in technical triplicate with viruses at a multiplicity of infection (MOI) of 0.03 plaque forming units (PFU)/cell. After 1 hr, cells were washed three times with FBS-free DMEM and cultured with FBS-free DMEM supplemented with 0.5μg/mL trypsin type IX. Viral supernatant and whole cell lysates were harvested at 1, 8, 16, 24, 32 and 48 hours post infection (hpi) and frozen at −80°C. Virus titres were determined by plaque assay. Whole cell lysates harvested in 350μL buffer RLT (QIAGEN) were used to determine the RNA levels by RT-qPCR and lysates harvested in 100μL 2X Laemmli buffer were used to analyse protein expression by western blotting, as described below.

### Viral RNA extraction and quantification by quantitative RT-PCR (RT-qPCR)

Primers for RT-qPCR targeting segments VP1 (SA11 and RF), VP4 (SA11) and NSP3 (RF) were designed using SSE v1.2 and OligoCalc software [70, 73] (Table 2).

RNA was extracted from infected cell lysates using RNeasy® Mini Kit (QIAGEN) with an on-column DNase treatment (QIAGEN) to remove possible DNA contamination according to manufacturer’s specification. The extracted RNA was dissolved in 20μL nuclease-free water (QIAGEN) and stored at −80°C.

To quantify total RNA levels of each segment relative to WT from timecourse experiments, one-step RT-qPCR was performed using SensiFAST™ SYBR® Lo-ROX One-Step Kit (Meridian Bioscience) with 2μL of RNA samples according to the manufacturer’s protocol on a Rotor-Gene Q apparatus (QIAGEN) using primers listed in Table 2. RT-qPCR cycling conditions were: 45°C for 10 min for reverse transcription, 2 min at 95°C, followed by 40 cycles of 10 sec at 95°C and 30 sec at 60°C. The conditions then increased from 50°C to 99°C at 1-degree increments to generate a melt curve to confirm specific amplification of each gene. For relative expression of each gene, the results were analysed according to the 2^-ΔCt^ method. Average Ct values of each gene were normalised to WT at 8 hpi (when RNA was first reliably detected for VP1, VP4 and NSP3) and the resulting ΔCt values were adjusted for primer efficiency. Three independent experiments were performed in technical duplicates.

To quantify the RNA copies of corresponding gene segments in mutant viruses for RNA:PFU ratios, RNA from cDNA (approximately 100ng) of each gene segment was synthesised using a MEGAscript™ T7 Transcription Kit (Invitrogen) followed by TURBO DNase treatment (Invitrogen) and a clean-up with MinElute PCR Purification Kit (QIAGEN) according to manufacturer’s instructions. The extracted RNA was dissolved in 10μL nuclease-free water (QIAGEN) and stored at −80°C. RNA was measured using Qubit™ RNA broad range assay kit (Invitrogen) and serial dilutions of RNA were used as standards in one-step RT-qPCR. A ratio of the transcript copy number of each gene segment in the mutant viruses to virus titre was calculated and normalised to WT. Three independent experiments were performed in technical triplicates with the standard done in duplicate.

The PCR efficiency (E) of each primer pair set was established by measuring serial dilutions of cDNA of each segment in triplicate and calculated based on the slope of the standard curve according to the formula E = (10^(−1/slope)-1^) x 100. Threshold cycle (Ct) values equivalent to mock samples and non-template control were considered to be negative.

### Western blotting (WB)

Whole cell lysates harvested in 2X Laemmli buffer from timecourse infections were boiled at 95°C for 10 min. Proteins were separated by SDS-PAGE on 10% or 4-20% Mini-PROTEAN® TGX™ Precast Protein Gels (Bio-Rad) according to the manufacturer’s instructions. Proteins were transferred onto a 0.2μm Cytiva™ Amersham Protran™ Nitrocellulose membrane (Fisher Scientific) and blocked with 2% horse serum in 1X Tris-buffered saline (20mM Tris, 150mM NaCl) containing 0.01% Tween® 20 detergent (Sigma-Aldrich) (TBS-T) for 1 hr at room temperature. Membranes were then probed with primary antibodies diluted in 2% horse serum/TBS-T and incubated overnight at 4°C with agitation (Table **4**). After three 5 min washes with TBS-T, membranes were incubated for 1 hr at room temperature with secondary antibodies diluted in TBS-T (Table **4**). Following three 5 min washes with TBS-T, membranes were imaged using an Odyssey ® XF imaging system (LI-COR). Analyses were performed with Image Studio™ Lite software (LI-COR).

**Table 4.**
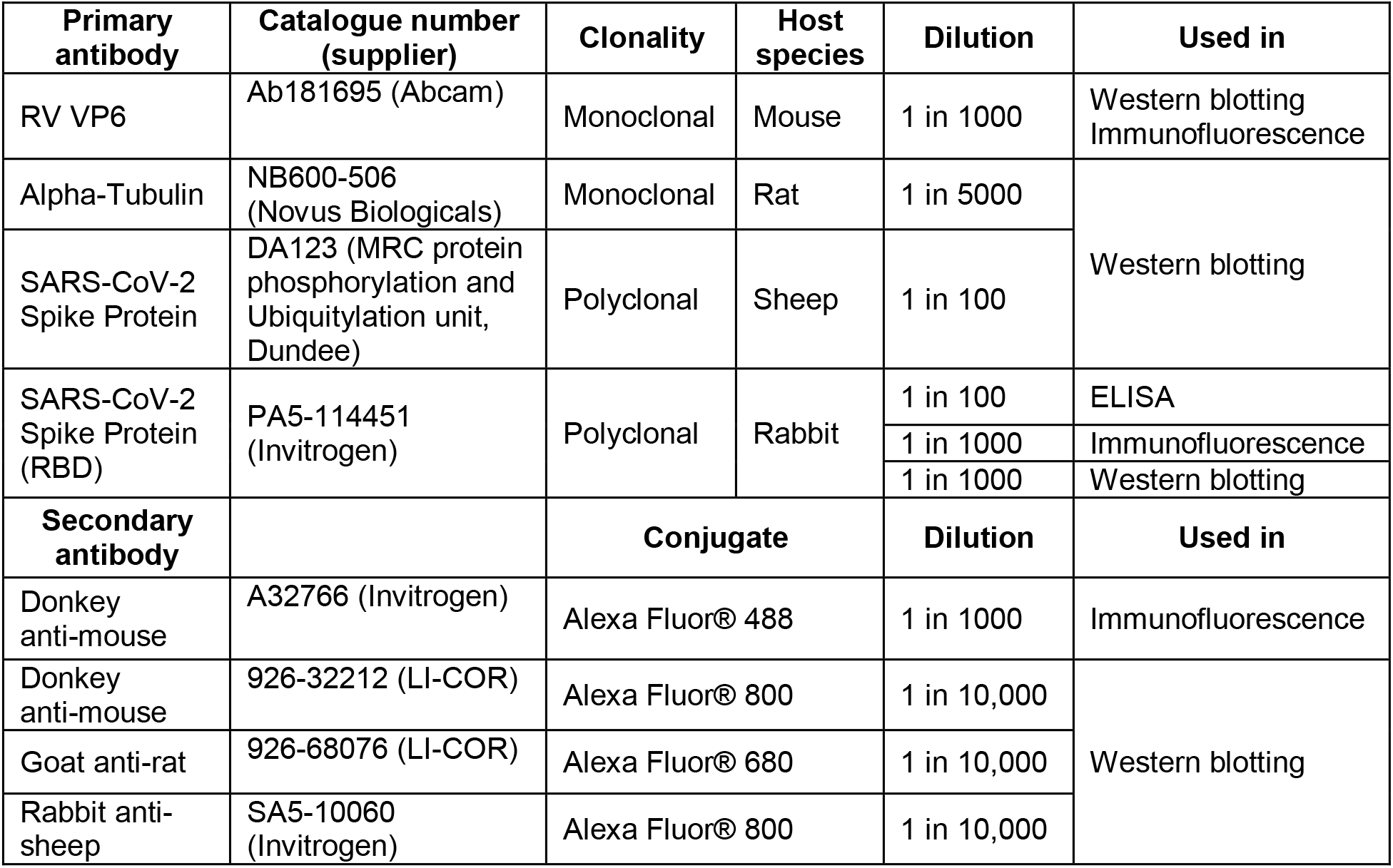
Primary and secondary antibodies used in this study.

| Primary antibody | Catalogue number (supplier) | Clonality | Host species | Dilution | Used in |
| --- | --- | --- | --- | --- | --- |
| RV VP6 | Ab181695 (Abcam) | Monoclonal | Mouse | 1 in 1000 | Western blotting<br>Immunofluorescence |
| Alpha-Tubulin | NB600-506 (Novus Biologicals) | Monoclonal | Rat | 1 in 5000 | Western blotting |
| SARS-CoV-2 Spike Protein | DA123 (MRC protein phosphorylation and Ubiquitylation unit, Dundee) | Polyclonal | Sheep | 1 in 100 |  |
| SARS-CoV-2 Spike Protein (RBD) | PA5-114451 (Invitrogen) | Polyclonal | Rabbit | 1 in 100 | ELISA |
|  |  |  |  | 1 in 1000 | Immunofluorescence |
|  |  |  |  | 1 in 1000 | Western blotting |
| Secondary antibody |  | Conjugate |  | Dilution | Used in |
| Donkey anti-mouse | A32766 (Invitrogen) | Alexa Fluor® 488 |  | 1 in 1000 | Immunofluorescence |
| Donkey anti-mouse | 926-32212 (LI-COR) | Alexa Fluor® 800 |  | 1 in 10,000 | Western blotting |
| Goat anti-rat | 926-68076 (LI-COR) | Alexa Fluor® 680 |  | 1 in 10,000 |  |
| Rabbit anti-sheep | SA5-10060 (Invitrogen) | Alexa Fluor® 800 |  | 1 in 10,000 |  |

### Virus purification and electrophoretic analysis of dsRNA and protein

Virions were semi-purified from SA11 and RF RV stocks using ultracentrifugation as described previously [71]. Briefly, for RNA visualisation, 50mL of clarified RV stock was pelleted through a buffered 25% (w/v) sucrose cushion (100mM NaCl, 10mM Tris-HCl [pH 7], 1mM EDTA) using a SW32Ti rotor in a Beckman Coulter Optima Max-E ultracentrifuge at 25,000 RPM for 90 min at 4°C. Viral pellets were resuspended in 350μL buffer RLT containing 3.5μL β-mercaptoethanol (Sigma-Aldrich) and dsRNA was extracted using an RNeasy® Mini Kit as described above. Purified RNA was treated with RNase A (Thermo Scientific) for 30 min at 37°C to specifically degrade single-stranded RNA and retain viral genomic dsRNA.

Extracted dsRNA was separated using 5% urea polyacrylamide gels in 1X Tris-borate-EDTA (TBE) buffer (89mM Tris-borate, 2mM EDTA [pH 8.3]). Gels were fixed in gel fixing solution (30% (v/v) methanol in 1X TBE buffer with 10% (v/v) acetic acid) and RNA was visualised using the Silver Stain Plus Kit (Bio-Rad) according to the manufacturer’s instructions. Gels were dried onto filter paper and imaged using a Samsung Xpress C480FW scanner.

To visualise viral proteins of VP4 mutants, 25mL of clarified RV stock was used for ultracentrifugation as described above. Viral pellets were resuspended in 1X TNC buffer (20mM Tris-HCl [pH 8], 100mM NaCl, 1mM CaCl_2_) and mixed 1:1 with 2X Laemmli buffer followed by SDS-PAGE using 4-20% Mini-PROTEAN® TGX™ Precast Protein Gels and staining with Coomassie Brilliant Blue R-250 (Bio-Rad) according to manufacturer’s instructions. Gels were imaged using the Samsung Xpress C480FW scanner. Quantification was performed by densitometry of scanned gel images using Image J. Values were corrected for background noise and normalised to those of the WT virus.

### Indirect-binding enzyme-linked immunosorbent assay (ELISA)

To determine the antigenicity of viruses expressing SARS-CoV-2 peptides, 200μL of either viral supernatant (VP4 mutants) or cell lysates (NSP3 mutants) diluted in 0.1M Carbonate-Bicarbonate Buffer (Sigma-Aldrich) was immobilized on clear 96-well ELISA plates (Greiner Bio-One) and incubated overnight at 4°C with agitation. Following two washes with PBS containing 0.05% Tween® 20 detergent (PBS-T) using a Biochrom Asys Atlantis microplate washer (Scientific Laboratory Supplies), plates were blocked with 2% horse serum/PBS at room temperature for 2 hr. After blocking, plates were incubated with 100μL/well primary antibody (Table **4**) diluted in 2% horse serum/PBS overnight at 4°C with agitation. Plates were then washed six times with PBS-T and incubated with horseradish peroxidase-conjugated goat anti-rabbit IgG (H + L) (1:2000) (Bio-Rad) diluted in 2% horse serum/PBS for 1 hr at room temperature with agitation. After 1 hr, plates were washed three times with PBS-T and incubated with 2,2′-Azino-bis(3-ethylbenzothiazoline-6-sulfonic acid) (Scientific Laboratory Supplies) substrate to allow the colour to develop after which the reaction was terminated by adding 70μL per/well of 1% SDS solution. The optical density (OD) was measured at 405nm using Cytation™ 3 Cell Imaging Multi-Mode Reader (Agilent) and data was analysed using BioTek Gen5 software (Agilent). Three independent experiments were performed with standards done in triplicate. For a positive control, an influenza A virus strain A/Puerto Rico/8/1934 was tagged with the same RBM sequence in the haemagglutinin protein (unpublished).

### Infection of bovine enteroids with NSP3 mutants

3D bovine enteroids were prepared as described in Hamilton *et al*., 2018 [74]. Infection of organoids was carried out as described by Derricott *et al*., 2019 [75] with modifications. 3D organoids were mechanically disrupted into multicellular fragments by pipetting to expose the apical surface of the cells in 80% of the enteroids. The sheared enteroids were counted using a bright-field microscope and diluted to the final concentration of 2500 enteroids/well. Each enteroid was estimated to contain approximately 40 cells. The sheared enteroids were then diluted to the appropriate concentration with IntestiCult™ Organoid Growth Medium (STEMCELL Technologies) supplemented with 10μM ROCK pathway inhibitor (Cayman chemicals), 10μM Galunisertib (Cayman chemicals) and 10μM p38 inhibitor (Enzo) and aliquoted into 15mL falcon tubes for each condition. The enteroid fragment suspension was infected with NSP3 mutants at an approximate MOI of 10 and incubated for 1 hr at 37°C. After 1 hr, the enteroid/virus suspension was centrifuged at 500 rpm for 2 min and washed five times by replacing the supernatant with equal volumes of fresh IntestiCult™. Enteroid pellets were then resuspended in the appropriate volume of IntestiCult™ and 500μL plated onto Corning® Matrigel® (Scientific Laboratory Supplies) coated coverslips in a pre-warmed 24-well plate.

### Immunofluorescent staining of bovine enteroids

Enteroids were fixed at 1 and 24 hpi with 4% paraformaldehyde for 1.5 hr at 4°C with agitation. Following three washes with PBS, enteroids were permeabilised with 0.5% Triton-X100 in PBS for 15 min and then blocked with 2% horse serum in PBS for 1 hr, all at room temperature. Enteroids were incubated with primary antibodies (Table **4**) overnight at 4°C with agitation, then washed three times with PBS and incubated with secondary antibodies (Table **4**) and phalloidin (F-actin detection) (1:100) (Invitrogen) for 1 hr at room temperature. All antibodies were diluted in 2% horse serum/PBS. Enteroids were then washed with PBS three times with the addition of 4’,6-diamidino-2-phenylindole (DAPI) nuclear stain (1:5000) (Invitrogen) for the final 10 min wash. Coverslips were rinsed in water and mounted onto microscope slides (Thermo Scientific) with ProLong™ Gold Antifade Mountant (Invitrogen), and imaged using a Zeiss LSM 710 confocal microscope at 630X magnification. Images were analysed using the Zen Black software and processed using Photoshop v23.1.1.

### Statistical analysis

GraphPad Prism v9 was used for all statistical analyses. Data are presented as mean and standard deviation from three independent experiments with technical duplicates unless otherwise stated. *P* values were determined by ratio paired t-test and were considered statistically significant at <0.05, unless data were normalised in which case paired t-tests were used.

## RESULTS

### *IN VITRO* SYNTHESIS OF RV PROTEINS TAGGED WITH SARS-COV-2 SPIKE PEPTIDES

We engineered a panel of SA11 strain VP4 plasmids with SARS-CoV-2 spike peptide sequences inserted into the hypervariable region, and a panel of RF strain NSP3 plasmids with 3’ tags of SARS-CoV-2 RBM or RBD with or without a separating T2A peptide (Fig. 1A). To confirm the expression of the spike epitopes by the mutated VP4 and NSP3 segments, coupled *in vitro* transcription and translation (IVT) reactions were carried out using rabbit reticulocyte lysate system supplemented with [^35^S] methionine (Fig. 1B and 1C).

Following SDS-PAGE and autoradiography, SA11 WT and mutated VP4 constructs produced polypeptide products of the expected protein size throughout (Fig. 1B). The WT RF NSP3 construct expressed a protein of expected size (Fig. 1C, lane 2). RF NSP3 segment incorporating the RBM motif produced a protein of a higher molecular weight consistent with its predicted size (Fig. 1C, lane 3). The T2A_RBM construct generated a product containing the NSP3 protein that ran slightly higher than WT NSP3 due to the residual C-terminal fusion of partial T2A sequence [76] (Fig. 1C, lane 4, red asterisk). The small size (~10kDa) of the unconjugated RBM peptide made it difficult to visualise due to co-migration with the dye front. Unseparated NSP3-T2A_RBM, seen as a minor product, was probably produced as a result of unsuccessful ribosome skipping at the T2A site [76] (Fig. 1C, lane 4, black asterisk). As expected, translation of NSP3 fused to the RBD peptide also produced a protein of a higher molecular weight than the WT (Fig. 1C, lane 5). For the T2A_RBD construct, untagged NSP3 was readily identifiable, and again a fainter band corresponding to the predicted molecular weight of fused NSP3-T2A_RBD was also seen (Fig. 1C, lane 6, red and black asterisks respectively). The RBD product (~23kDa) was reproducibly not detectable, possibly due to discontinued translation as a result of ribosome fall-off at the T2A site, or degradation in the rabbit reticulocyte lysate system. These results show that the spike epitopes were successfully translated in a cell-free system and that the T2A peptide was functional.

### RESCUE OF VP4 AND NSP3 MUTANT VIRUSES

To generate the VP4 and NSP3 mutant viruses, we used an adapted version of the previously published SA11 reverse genetics system or our established plasmid-only RF reverse genetics system respectively [42, 43] (Fig. 2A). With the exception of the NTD mutant, all VP4 mutants displayed significantly lower titres compared to the WT, with approximately two-log_10_ decrease in the titres observed across the panel (Fig. 2B). Nonetheless, all VP4 mutants were successfully rescued on all attempts. The VP4 mutants exhibited a similar but smaller speckled plaque phenotype to the WT (Fig. 2C). Statistical analyses of plaque sizes were not performed due to the ambiguities in determining the peripheries of individual plaques and their non-uniform diameters within a well.

**Figure 2.**
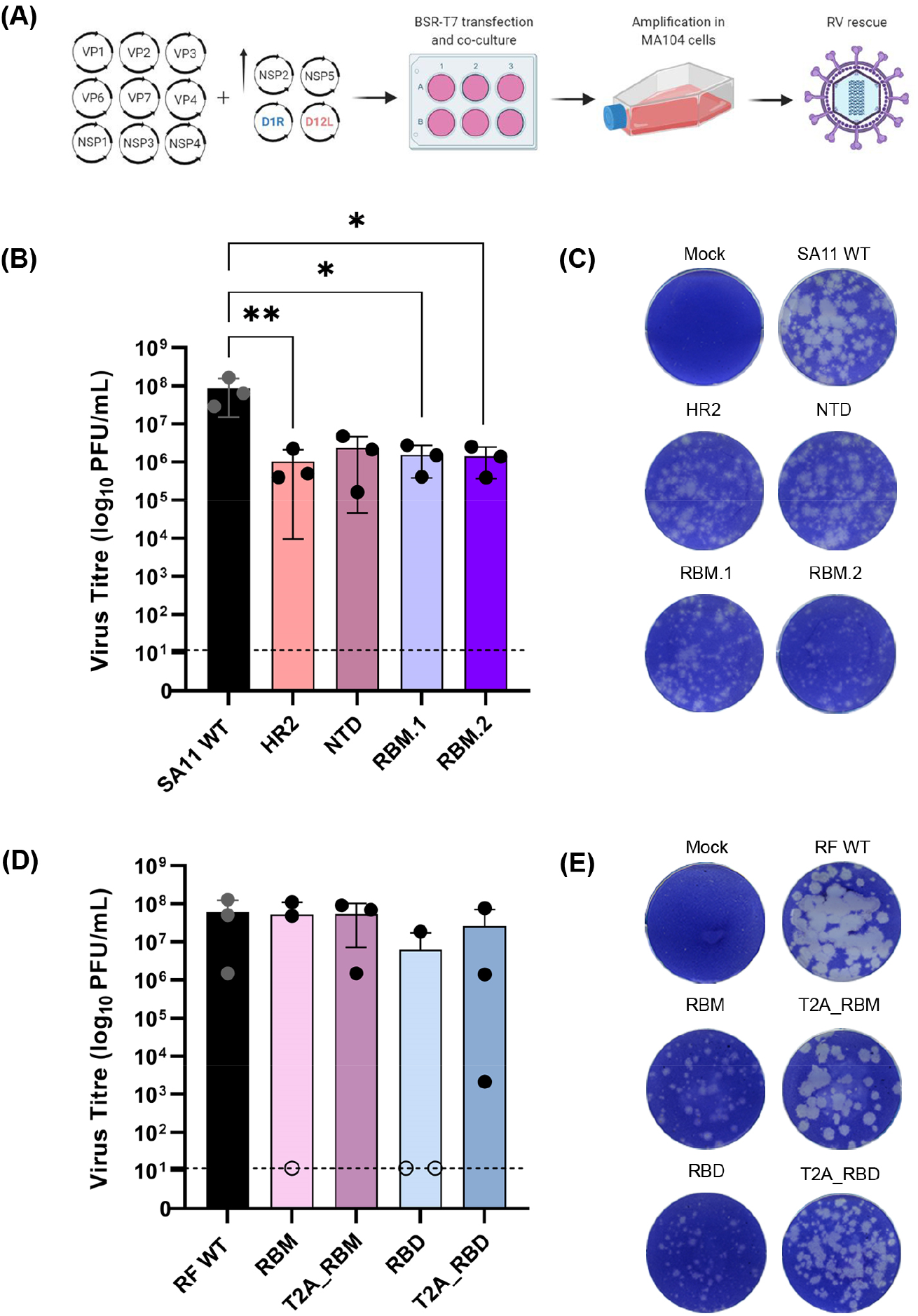
Generation of VP4 and NSP3 mutant viruses using plasmid-only based reverse genetics system. **(A)** Schematic representation of a 13-plasmid system for generation of mutant viruses expressing spike epitopes (*Adapted from Kanai et al., 2017 and Komoto et al., 2018; created with BioRender.com*) [42, 43]. Full length cDNAs representing each of the 11 gene segments were transfected into BSR-T7 cells with increasing amounts of two plasmids carrying NSP2 and NSP5 genes, along with two plasmids expressing vaccinia virus capping enzyme genes (D1R and D12L). Mutant viruses were rescued following amplification in MA104 cells. Viral titres and plaque morphology versus WT for VP4 **(B – C)** and NSP3 **(D – E)** mutants. Dashed line represents detection threshold. **(B)** \**p ≤ 0.05* (RBM.1 *p = 0.0205*; RBM.2 *p = 0.0217*), \*\**p ≤ 0.01* (HR2 *p = 0.0018*). Representative results from three independent rescues is shown (except for RBD, which only rescued once). In **(D)** open circles show failed rescues plotted at the limit of detection.

In contrast, compared to the WT virus, there were no statistically significant differences observed in titres of any of the NSP3 mutants (Fig. 2D). However, out of three attempts, RBM was successfully rescued twice whilst T2A_RBM was rescued at all times, with one of the rescues showing a two-log_10_ lower titre (Fig. 2D). The RBD mutant was only rescued once with a titre that was a log_10_ lower than the WT (Fig. 2D). Although T2A_RBD rescued on all attempts, it delivered a five-log_10_ variation in titres between the rescues (Fig. 2D). The viruses with NSP3 tags displayed smaller plaques than the WT, except T2A_RBM, whose plaques had a similar morphology to that of WT (Fig. 2E). Notably, in the presence of T2A peptide, plaque sizes were larger than when RBM or RBD was fused directly to the C-terminus of NSP3 without T2A (Fig. 2E).

Thus, small peptide insertions into the hypervariable region of SA11 VP4 significantly reduced the virus titre, while the efficiency of viral rescue was affected by the size of the peptides fused to the C-terminus of RF NSP3. Furthermore, these data demonstrated that the RF strain of RV was successfully rescued from cloned cDNAs (confirmed by sequencing, data not shown).

### CHARACTERISTICS OF VP4 AND NSP3 MUTANT VIRUSES

Following virus rescue, we first examined the effects of peptide insertion on RV replication in cell culture. Multi-step growth curves were performed for the VP4 and NSP3 mutants after infection of MA104 cells at a low MOI. From 24 hpi, all VP4 mutants had significantly lower titres (≥ one-log_10_) than the WT virus for at least one timepoint during the infection (Fig. 3A). Next, we assessed the impact of the introduced mutations on the total RNA expression levels for VP1 and VP4 viral transcripts. No marked differences in transcript levels of either VP1 (Fig. 3B) or VP4 (Fig. 3C) were observed (*p ≥ 0.05*, paired t-test).

**Figure 3.**
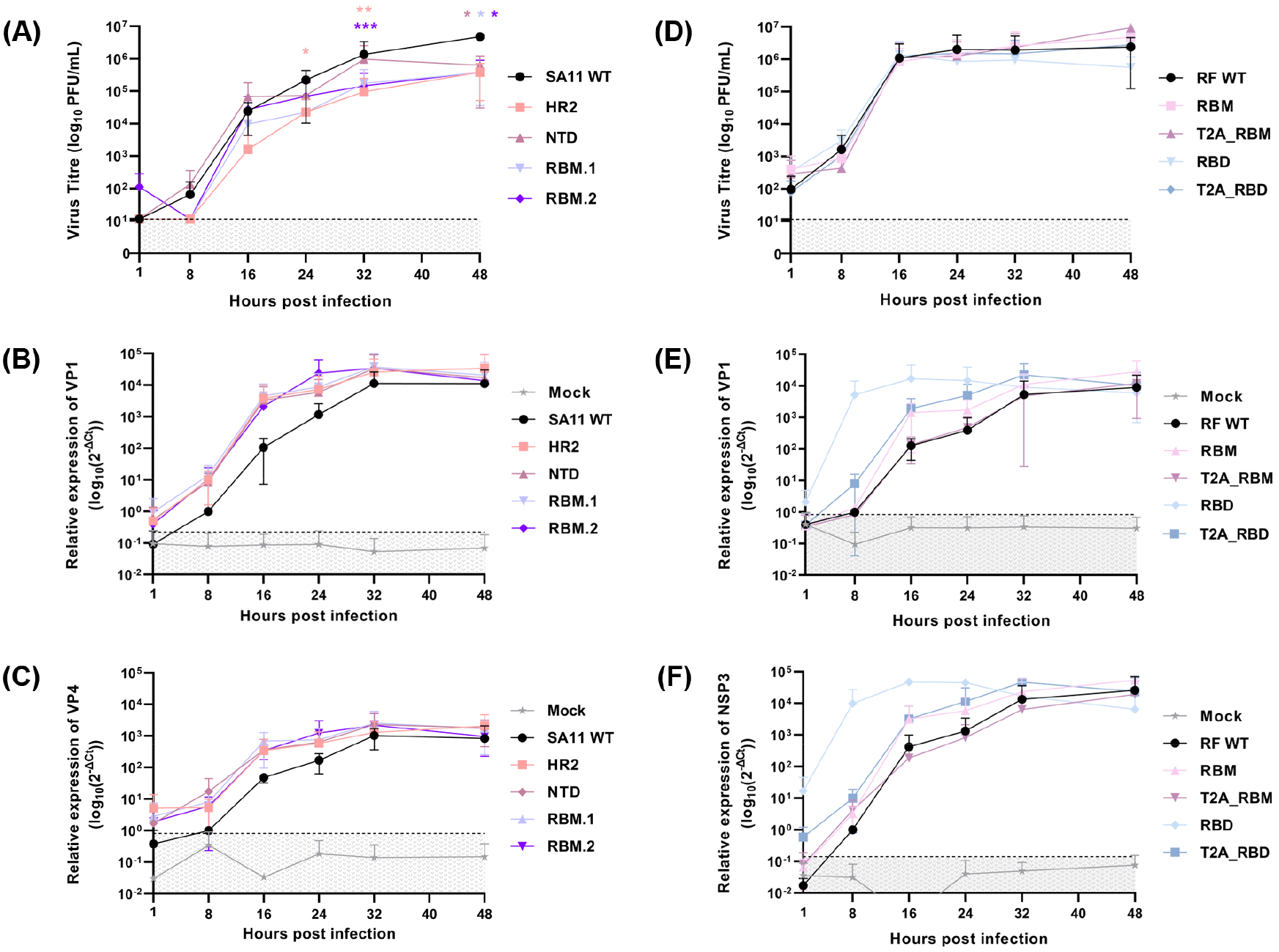
Replication kinetics and viral RNA content of WT and mutant viruses. **(A** and **B)** Multi-step growth curves for VP4 and NSP3 mutant viruses respectively. **(A)** \**p ≤ 0.05* (24 hpi HR2 *p= 0.041*; 48 hpi NTD *p = 0.047*, RBM.1 *p = 0.045*, RBM.2 *p =0.031*), \*\**p ≤ 0.01* (32 hpi HR2 *p = 0.009*), \*\*\**p ≤ 0.001* (32 hpi RBM.2 *p = 0.0006*). **(C – F)** Total cellular RNA levels of VP1, and VP4 (SA11 panel) or NSP3 (RF panel) transcripts. Data are from three independent experiments. Dashed line represents detection threshold.

Conversely, the NSP3 mutants followed similar replication kinetics to RF WT with three-log_10_ increases in titres between 8 and 16 hpi, with titres plateauing thereafter (Fig. 3D). Throughout, no major differences in titres between any of the viruses in the panel were observed. For the NSP3 virus panel, mutants produced higher levels of VP1 (Fig. 3E) and NSP3 (Fig. 3F) transcripts than the WT virus earlier in the time course. Higher RNA levels of VP1 and NSP3 in the RBD mutant is likely a result of higher RNA input (Fig. 3E–F respectively). Neither VP1 nor NSP3 transcript levels differed significantly throughout the time course (*p ≥ 0.05*, paired t-test) (Fig. 3E–F).

Western blot analyses were then used to characterise the production of SARS-CoV-2 spike polypeptides and RV VP6 using whole-cell lysates from the infection time course. As different SARS-CoV-2 peptides were introduced in each mutant, anti-spike antibody affinity may vary between mutants and so expression levels cannot be compared. Due to lack of available antibodies, we were unable to measure RV VP4 and NSP3 protein levels directly.

For the detection of various SARS-CoV-2 spike peptides in cells infected with the VP4 mutants, a polyclonal SARS-CoV-2 spike antibody was used. As the spike peptides were introduced into the hypervariable region of VP4, we considered the possibility of detecting the full length VP4 (sizes indicated in Table 3), and/or the VP8* cleaved product containing the spike peptides. At 32 hpi, full length VP4 product was detected in HR2, NTD and RBM.2 mutants (Fig. 4A, black asterisks). No SARS-CoV-2 spike peptide signal was detected for the RBM.1 mutant, likely reflecting selective clonality of the antibody used (Fig. 4A). Interestingly, at 32 hpi, the upper band representing the uncleaved VP4 product was brighter for RBM.2 (Fig. 4A, black asterisks), whereas for NTD, the lower band possibly representing the VP5* cleaved product was stronger than the upper band (Fig. 4A, green asterisks). These differences could be due to the distinct efficiencies of VP4 processing caused by the inserted peptides. The hypervariable region where the spike peptides were introduced is within the VP8* domain, and we would expect a product of around 30-31kDa in the event of VP4 cleavage. Possibly, this is evident in NTD at 32 and 48 hpi (Fig. 4A, blue asterisks), although the NTD mutant showed several unexpected bands and so this may reflect non-specific antibody binding; a faint product of similar molecular weight was also detected in the mock samples at 32 hpi (Fig. 4A).

**Figure 4.**
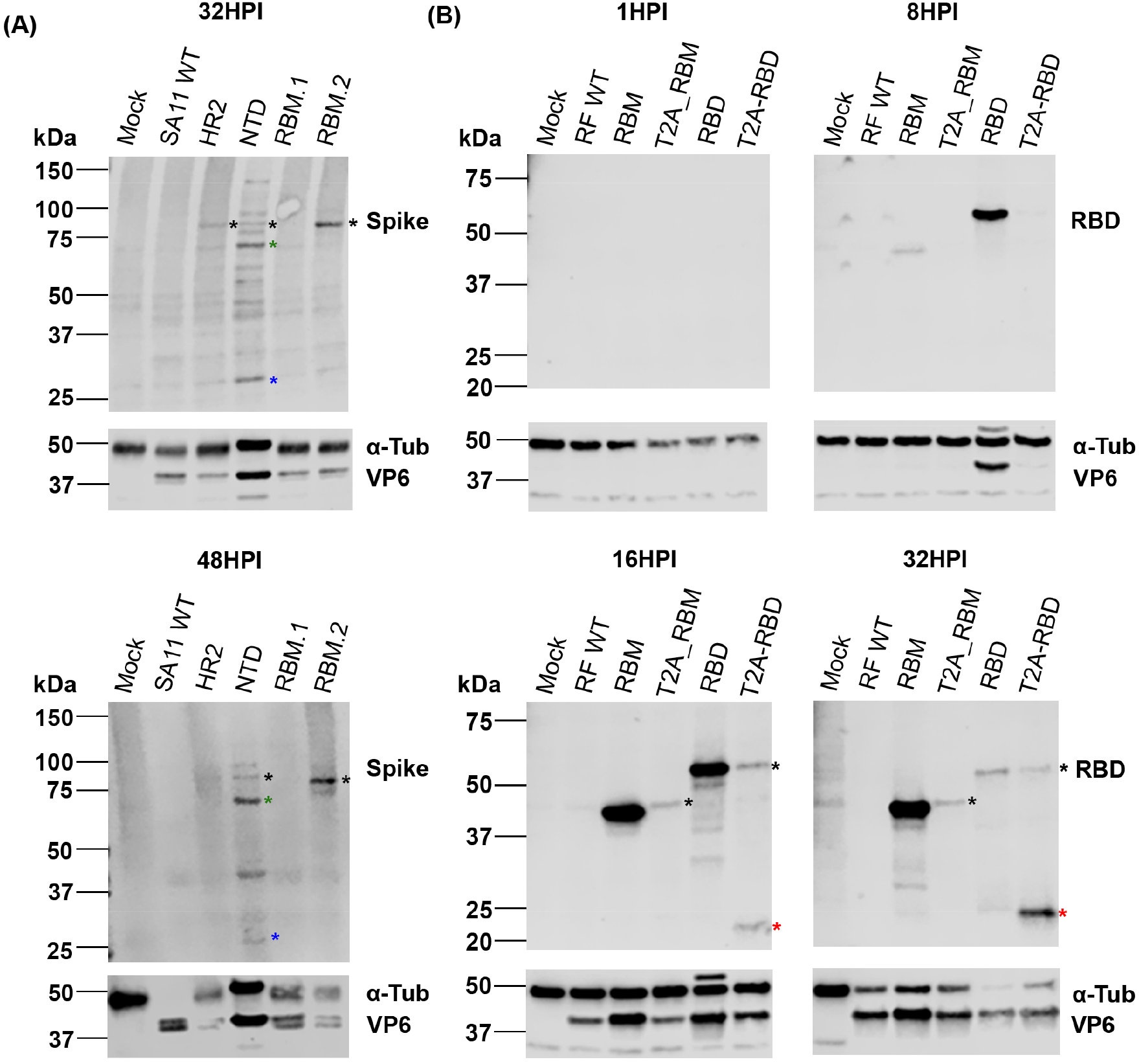
Expression of VP6 and SARS-CoV-2 spike peptides in infected cells. Cells infected at low MOI were harvested at 1, 8, 16, 24, 32 and 48 hpi. Whole-cell lysates were analysed by SDS-PAGE and western blot using polyclonal antibodies against RV VP6, and spike for SA11 mutants **(A)** or RBD for RF mutants **(B)**. Alpha-tubulin (α-Tub) was used as a loading control. In **(A)** black asterisks denote uncleaved VP4 product. Cleaved VP4 products VP5* and VP8* are marked by green and blue asterisks respectively. In **(B)** black asterisks show T2A read-through product and red asterisks identify separated products. Representative results from three independent experiments are shown. Position of molecular weight markers are indicated (kDa).

We also examined RV VP6 production which was detected in all VP4 viruses at 32 hpi (Fig. 4A). VP6 signal intensity did not correlate with spike signal intensity across viruses, again likely due to variable affinities of the spike antibody for the different spike peptides incorporated into VP4 (Fig. 4A).

In cells infected with NSP3 mutants, detection of both RBD and RBM peptides was possible using a polyclonal SARS-CoV-2 RBD antibody. Cross-reactivity for the RBM–expressing virus was first detected at 8 hpi, with increased levels present at 16 and 32 hpi (Fig. 4B). Antibody cross-reactivity for the T2A_RBM virus was only visible between 16 and 32 hpi (Fig. 4B). The band detected is consistent in size with an unseparated NSP3-T2A_RBM protein product (Table 3, Fig. 4B, black asterisks). The RBM peptide of around 10kDa was not detected, consistent with its *in vitro* translation efficiency (Fig. 4B, Fig. 1B). For the RBD virus, expression of the RBD peptide was first observed at 8 hpi but declined from 16 hpi paralleling the disappearance of tubulin (likely due to cell death) (Fig. 4B). In the T2A_RBD mutant, both NSP3-conjugated and unconjugated RBD products were detectable from 16 hpi (Fig. 4B, black and red asterisks respectively). Over time, the RBD peptide (~23kDa) became more apparent, as the NSP3-RBD signal diminished, confirming the functionality of the T2A element (Fig. 4B, red asterisks).

It is unclear why the T2A-induced ribosomal skipping appeared to improve in efficiency over the course of infection. It is possible that the stability of the fused peptides is lower than the separated peptides. Similarly, over the course of infection, RBM protein levels increased throughout, whereas RBD protein levels increased until 16 hpi after which they dropped dramatically (Fig. 4B).

The expression of VP6 was first observed at 8 hpi during infection with the RBD mutant, correlating with the signal of the RBD peptide, but was only detected for the remaining viruses from 16 hpi (Fig. 4B). Since equal MOIs were used for time course infections, higher input of genomic RNA copies could explain earlier V6 detection in the RBD mutant. This is consistent with the higher transcript levels detected for the RBD mutant at 8 hpi (Fig. 3E–F). In the event of a packaging defect of the RBD mutant more input genome copies would be required to deliver equal numbers of infectious particles.

Infection with both mutants resulted in a similar drop in tubulin levels (Fig. 4A–B), suggesting that the two mutants have different protein turnover rates.

In summary, introducing short SARS-CoV-2 peptides into the hypervariable region of VP4 impacted the virus yield, whereas fusing SARS-CoV-2 spike peptides to the C-terminus of NSP3 with or without T2A did not affect the virus titre or replication kinetics. Nevertheless, SARS-CoV-2 spike peptides were detectable in the majority of mutants using polyclonal antibodies, encouraging further investigations into the potential of these strategies for heterologous peptide presentation.

### EFFECT OF GENE MUTATION ON VIRAL GENOME PACKAGING

For viruses of the *Reoviridae* family, genome packaging is a tightly orchestrated and controlled process, so increasing the RV genome size may affect packaging efficiency [30, 77–79]. To examine whether our mutations may have affected genome packaging, RNA was extracted from equal volume of purified viruses and analysed by urea-PAGE and silver staining.

The VP4 virus panel showed the expected constellation of genome segments, but as samples were resolved for an extended period of time to visualise the small changes in segment 4 (VP4) band sizes expected as a result of SARS-CoV-2 spike peptide insertion, segments 10 and 11 ran off the gel (Fig. 5A). When gels were run for a shorter time, no differences in band densities of segments 10 and 11 were seen (data not shown). Mutated VP4 segments migrated more slowly than the WT VP4 segment, reflecting the various sizes of inserted spike sequences (Fig. 5A, highlighted in red). Densitometry analysis showed no substantial differences in band density between mutated VP4 and WT VP4 segments (normalised to WT: HR2 = 0.96, NTD = 1.11, RBM.1 = 0.87, RBM.2 = 0.96), suggesting no obvious packaging defects were introduced by small sequence insertions into the hypervariable region of VP4 (Fig. 5A).

**Figure 5.**
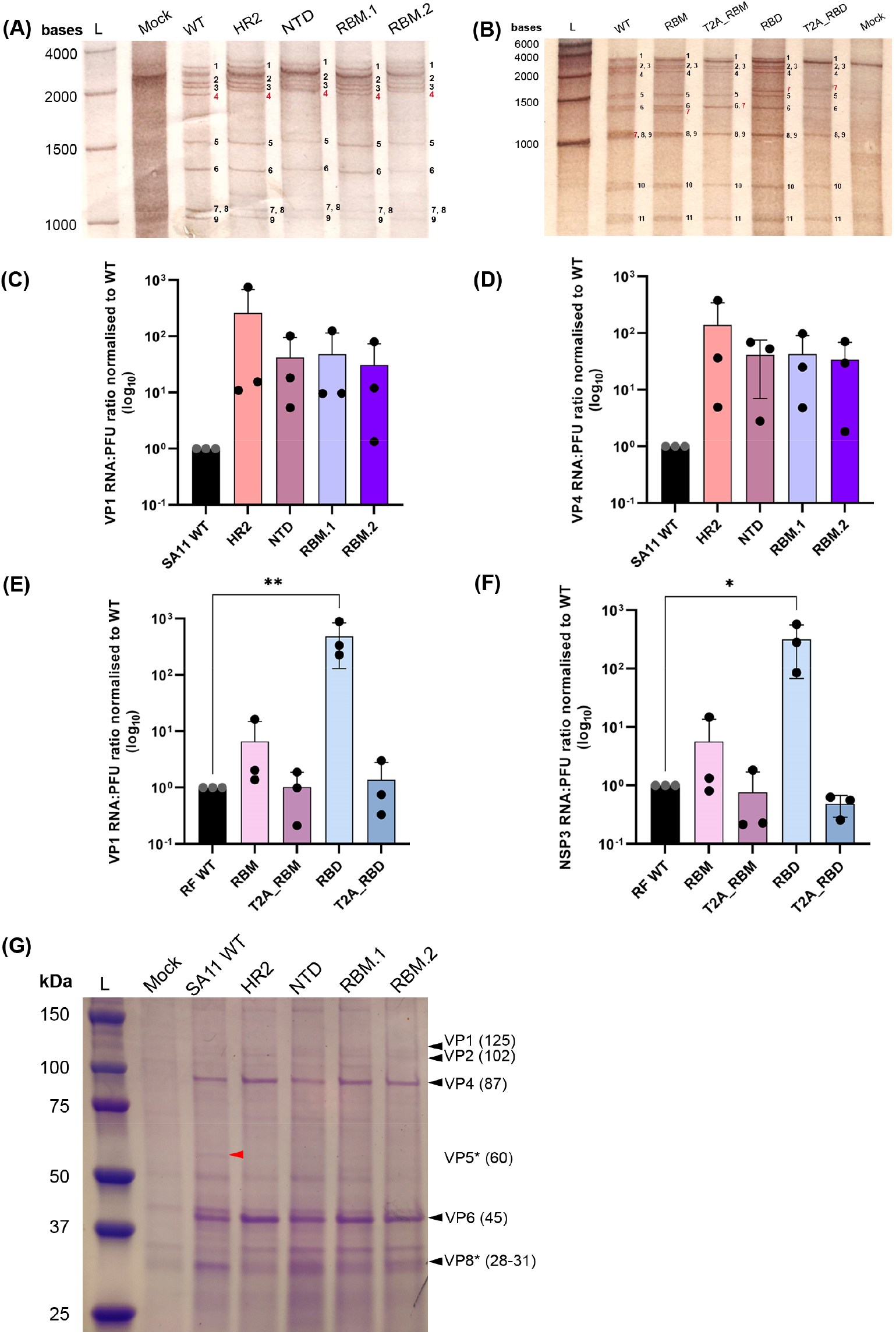
Impact of VP4 and NP3 mutation on packaging of RV RNA and proteins. Extracted RNA from virus stocks was analysed by urea-PAGE and silver staining for SA11 mutants **(A)** and RF mutants **(B)**. Individual RNA segments are labelled in black, and mutated RNA segment notations are in red. Lane “L” represents High Range RNA Ladder showing band size of RNA transcripts. RNA:PFU ratios of VP1 and either SA11 VP4 **(C – D)** or RF NSP3 **(E – F)** genes were determined by RT-qPCR and a ratio of copy number to viral titre was calculated; values were then normalised to WT. Dots represent individual samples from three independent rescues with each performed in triplicate. **(E)** \*\**p ≤ 0.01* (RBD *p = 0.005*). **(F)** \**p ≤ 0.05* (RBD *p = 0.010*). **(G)** Electrophoretic profile of viral proteins from purified VP4 mutants visualised using Coomassie brilliant blue. Lane “L” represents protein ladder indicating the molecular weight markers (kDa). Black triangles show the positions of viral proteins and their expected protein sizes (in brackets) are indicated (kDa). VP3 and VP7 proteins were not detected. VP5* protein was detected in WT only (red triangle).

The resolved segments for the NSP3 virus panel also showed the expected pattern of 11 RNA segments, as well as a prominent background band present in a mock infected sample that migrated between segments 1 and 2 (Fig. 5B). Segment 7 (encoding NSP3) from all mutants migrated notably slower than from WT virus, corresponding with its increased gene size (Fig. 5B, highlighted in red). Co-migration of segments 7, 8 and 9 in the WT made it difficult to reliably separate segment 7, precluding direct quantitative analyses (Fig. 5B). Nevertheless, no obvious defect in packaging was observed through visualisation of the complete genome of RF mutants.

To further evaluate whether virion infectivity may have been affected by genome mutagenesis, the genome copy number to PFU ratio was determined for the two panels of viruses, measuring VP1 and either VP4 (SA11 panel) or NSP3 (RF panel) segments (Fig. 5C–F). No significant differences were observed in the levels of VP1 and VP4 segments across the VP4 mutants relative to WT, although for all mutants a 1-2 log_10_ increase in RNA copies required to make an infectious virion was observed for both VP1 and VP4 (*p ≥ 0.05*, paired t-test) (Fig. 5C–D).

In contrast, all NSP3 mutants, with the exception of RBD, had equivalent numbers of VP1 and NSP3 segments (Fig. 5E–F). RBD had a significantly higher VP1 (*p = 0.005*, paired t-test) and NSP3 (*p = 0.010*, paired t-test) segment copy number:PFU ratio, with over 100-fold more copies of RNA required to make a fully infectious particle (Fig. 5C–D).

VP4 is important for viral attachment and entry as well as for the maturation of TLPs that constitute an infectious virus [1]. Therefore, it was considered that mutation of VP4 may affect the assembly of the viral structural proteins. To test this, purified VP4 viruses were further analysed by SDS-PAGE followed by Coomassie staining (NSP3 viruses were not included because the spike epitopes were fused to NSP3 which is not incorporated into virions) (Fig. 5G). Apart from VP3 and VP7 proteins, major structural proteins were detected throughout (Fig. 5G). VP4 was readily detectable for all viruses in the panel and so amounts were quantified by densitometry and normalised to that of WT virus. This showed no obvious VP4 incorporation defects, with all mutant viruses showing very similar VP4 amounts to the WT virus (Relative to WT, values were: HR2 = 1.07, NTD = 0.99, RBM.1 = 1.09, RBM.2 = 1.06). Thus, the mutant VP4 proteins appear to be incorporated into virus particles with unaltered efficiency.

A notable difference between the WT virus and its mutants was the cleavage pattern of VP4 in the presence of trypsin used for propagating the panel of mutant viruses. A band of the predicted size for VP5* was visible in the WT (Fig. 5G, red arrow). However, no corresponding band was visible for any of the VP4 mutants expressing heterologous peptides, further supporting our earlier hypothesis that peptide insertion into the VP4 hypervariable region impairs its processing into VP8* and VP5* (Fig. 4A).

In contrast, VP8* was detected in all mutants, and it migrated slightly higher than WT VP8, corresponding to the additional peptide sequences present in the hypervariable domain of VP4 (Fig. 5G). Nevertheless, VP8* band density was stronger for WT than for the mutants, again consistent with inefficient tryptic cleavage of VP4.

Overall, no major effect on viral assembly was observed in either of the mutant viral panels, with the exception of inserting the large RBD tag into NSP3 which caused a defect that could be overcome by incorporation of a T2A element.

### DETECTION OF SARS-COV-2 SPIKE ANTIGENS BY INDIRECT ELISA

To further investigate the viability of using RV as a delivery vector for SARS-CoV-2 spike antigens, we assessed whether the RV mutants expressing spike peptides would cross-react with RBD antibodies in an indirect ELISA. For the VP4 mutants, no antibody cross-reactivity was identified in this assay using the SARS-CoV-2 RBD antibody (Fig. 6A). Additionally, no cross-reactivity was observed using the SARS-CoV-2 spike antibodies to target the various spike peptides in the VP4 mutants, or with the PR8-RBM positive control (data not shown).

**Figure 6.**
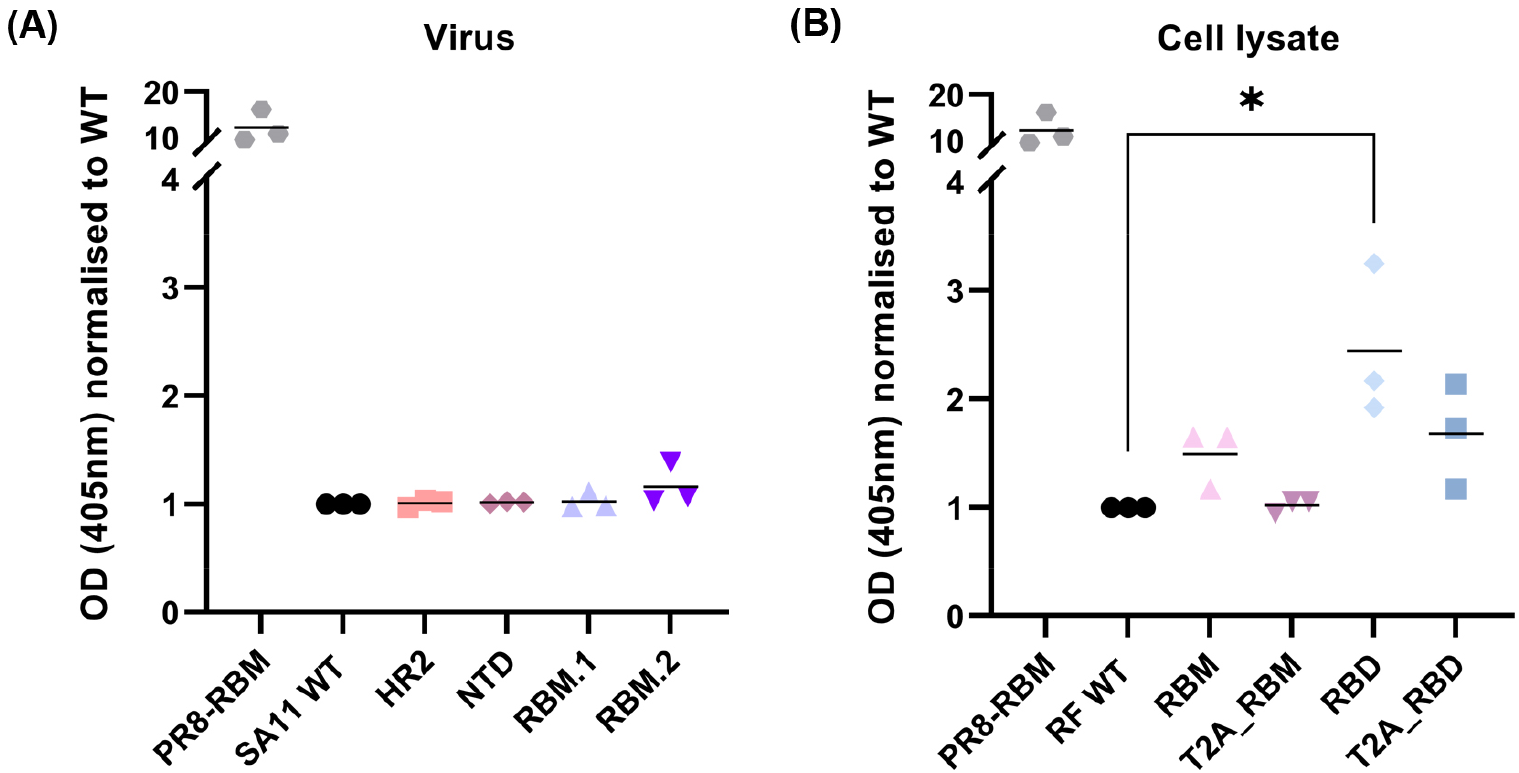
Detection of spike antigens expressed by mutant viruses. Presence of spike antigens were evaluated by indirect ELISA using viral supernatant of VP4 mutants **(A)** and cell lysates for NSP3 mutants **(B)**. Coloured symbols represent individual data points obtained from three independent experiments. OD (405nm) signal for all mutants was normalised to WT (background signal). PR8-RBM represents positive control. **(A)** *p > 0.05*. **(B)** \**p ≤ 0.05* (RBD *p = 0.032*).

Except the T2A_RBM mutant, all NSP3 mutants consistently showed a higher signal than the background signal from the WT virus, although only the RBD mutant showed a significantly higher signal than WT (*p = 0.032*, paired t-test) (Fig. 6B).

Based on these observations, RF NSP3 may be a better target for heterologous peptide conjugation than SA11 VP4 for expression of immunogenic antigens, which is at least partly attributable to the larger insertion site tolerated in the corresponding genome region.

### BOVINE ENTEROIDS AS A SPECIES-SPECIFIC MODEL FOR VIRAL INFECTION

Cell culture-based assays identified the RF NSP3 gene as a strong candidate for expressing immunogenic heterologous peptides. Orally administered live attenuated RV vaccines replicate in the gastrointestinal tract, and so bovine intestinal organoids containing enterocytes, goblet, Paneth, enteroendocrine and stem cells [75] were used to investigate if a more physiologically representative system was susceptible to infection with the NSP3 mutants. A bovine organoid system was chosen as the RF strain was first identified in diarrhoeic calves [80].

3D bovine enteroids were infected with NSP3 mutants at an approximate MOI of 10, stained with anti-VP6 and anti-RBD antibodies and imaged by confocal microscopy (Fig. 7A and 7B respectively). At 24 hpi, VP6 was predominantly detected in the epithelium comprised of mature enterocytes lining the apical surface of the organoid lumen (Fig. 7A), which is consistent with previous findings showing their preferential infection by RVs [19, 20]. In contrast, VP6 distribution in RBM and T2A_RBM mutants was detected around the nuclei of cells located within the organoid lumen and in the punctate cytoplasmic inclusion bodies, most likely viroplasms, sites of RV replication and assembly [1] (Fig. 7A, VP6 panel). VP6 signal in the RBD and T2A_RBD mutants was not apparent in the lumen and was mainly detected around the nuclei of cells in the organoid lining (Fig. 7A, VP6 panel).

**Figure 7.**
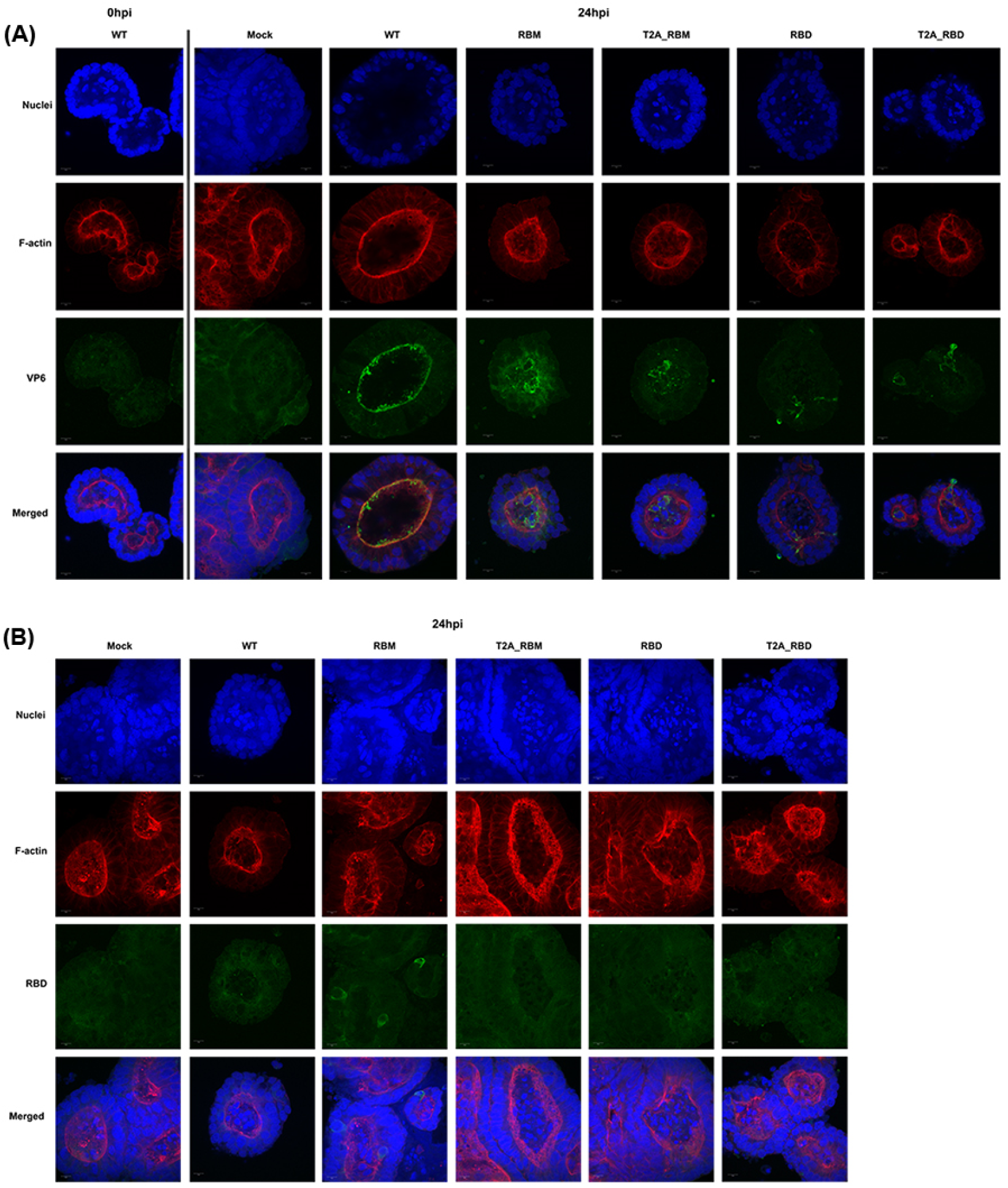
Bovine enteroids infected with NSP3 mutant viruses. 3D bovine enteroids were infected with the panel of NSP3 mutants, or mock at an approximate MOI of 10 PFU per cell and fixed at 24hpi, or at 0 hpi for WT as a control for background signal. Cells were stained for RV VP6 **(A)** and RBD of SARS-CoV-2 spike **(B)**. Scale bar represents 10μm.

Following staining with the SARS-CoV-2 RBD antibody, RBM signal was detected around the nuclei of cells in the organoid lining while other mutants did not produce any visible anti-RBD cross-reactivity (Fig. 7B). This is comparable to western blotting data where RBM also gave the strongest signal (Fig. 4B).

## DISCUSSION

We have used simian SA11 and bovine RF RV strains as viral vectors to express various SARS-CoV-2 spike epitopes as a model system with which to test the potential for expression of multivalent antigens. Tagging of the VP4 protein found on the surface of the virion with smaller peptides consistently impaired viral growth and did not yield strong antibody cross-reactivity. Conversely, relatively large foreign sequences could be tagged to the C-terminus of NSP3 protein, mostly without impairing viral titres and replication kinetics, and cross-reactivity with SARS-CoV-2 RBD antibodies was also demonstrable.

To investigate the feasibility of using RV as an expression vector, we analysed the effect of introducing SARS-CoV-2 spike peptides into the ‘head’ region of the VP4 (i.e. VP8* domain), outside the sialic acid binding domain. Since VP8* is used in several vaccine platforms such as protein subunit or nanoparticle vaccines to induce RV-specific neutralising antibodies [81–86], we hypothesised that the expression of SARS-CoV-2 spike peptides by VP8* may similarly generate neutralising antibodies. However, incorporation of various spike epitopes into the VP8* lectin domain between amino acid positions 164 – 198 consistently showed significant decrease in viral titres, despite having no obvious effect on the rescue efficiency (Fig. 2B).

It is possible that reduced infectivity of VP4 mutants could be due to inefficient conformational transition of VP4 following proteolytic cleavage [1, 23, 87]. Following cell attachment, VP8* lectin domains at the tip of VP4 dissociate and expose the hydrophobic loops of the VP5* β-barrel domains [21, 67, 68]. This enables interaction of the VP5* hydrophobic loops with the lipid bilayer and perforation of the target membrane by the VP5* foot domain, leading to RV entry, analogous to refolding of influenza virus haemagglutinin during membrane fusion [67, 68, 88–93]. Therefore, our mutation of the VP8* lectin domain may have indirectly affected the association of the hydrophobic loops of the VP5* β-barrel domains with the target membrane.

Alternatively, while we saw no obvious effects of VP4 mutagenesis on genome packaging (Fig. 5A, C–D), the absence of the VP5* detection in the mutants (where a band of the correct size was identified in the WT virus) could indicate inefficient cleavage of VP4 on TLPs (Fig. 5G). *In vitro* assembly of TLPs has shown that DLPs require addition of VP4 before VP7, and that the tryptic cleavage occurs following addition of VP7 [90, 94]. Interaction of VP7 subunits with the VP5* foot domain stabilise and anchor the VP4 onto the virion, allowing the protease to cleave only the linker sequence that bridges VP8* and VP5*, which in our mutants is intact [22, 23, 67, 68]. It is possible that our peptide insertions may have altered the VP4 conformation at this interaction site, disrupting VP4 cleavage and rendering VP5* undetectable in our assays (Fig. 4A and 5G).

Finally, we were unable to detect any cross-reactivity of the SARS-CoV-2 spike peptides with spike (not shown) or RBD antibodies in ELISA (Fig. 6A), further suggesting inefficiency of using SA11 VP4 as an expression vector for foreign peptides, at least using the chosen peptides. It is possible, however, that other peptides may be immunogenic when presented this way, since VP4 mutants were detectable in western blots.

Previous studies have shown that SA11 NSP3 is able to tolerate insertions of foreign sequences at its C-terminus [45, 51, 52]; we have demonstrated that this is also possible using the bovine RF strain NSP3. This is advantageous as the current live attenuated pentavalent RV vaccine, RotaTeq, utilises a bovine RV backbone reassorted with different human strains and the use of a non-primate origin RV vector helps to circumvent pre-existing immunity [10, 69, 95].

The RF strain of RVs demonstrates a positive feedback mechanism for NSP3 mRNA translation where the 3’ GACC end of mRNA molecules and both domains (N- and C-terminal) of NSP3 are required [96]. Accumulation of the first NSP3 molecules generated during infection triggers NSP3-dependent translation by specifically binding the terminal 3’ GACC of mRNA, and once established, NSP3 is available for translation of other viral mRNAs [96]. Depending on the virus strain, the 3’ GACC canonical sequence differs in some segments of RV, and the same positive feedback mechanism is not observed in the SA11 strain [1, 96]. The finding that RF strain NSP3 can tolerate large C-terminal tags was therefore not necessarily expected.

Rescue of the RF mutant with the RBD peptide (193 amino acids) fused directly to NSP3 was achieved only once in three attempts (Fig. 2D), possibly reflecting an impaired function of NSP3. We also found that the RNA:PFU ratio was affected in the RBD mutant only (Fig. 5E–F), and that copy numbers of NSP3 and VP1 encoding transcripts were both affected. This cannot be attributed to the longer length of this segment, as the NSP3-T2A_RBD mutant did not demonstrate any packaging defect (Fig. 5E–F). Therefore, NSP3 exerts its role in virus replication by regulating viral mRNA translation. This is consistent with the observed increase in RNA:PFU ratios being nonspecific to a particular segment.

With the exception of T2A_RBM, NSP3 mutants expressing SARS-CoV-2 spike peptides cross-reacted with the SARS-CoV-2 RBD antibody (Fig. 6B), supporting the previously proposed viability of using NSP3 as a tagging system [52] and demonstrating its application to other RV strains. Follow up studies using sera from COVID-19 patients are needed to confirm antibody responses to the antigens produced by the NSP3 mutants.

Overall, we conclude that the diminished infectivity of VP4 mutants accommodating only small peptide insertions further limits the use of the hypervariable region for foreign sequence expression and thus VP4 as a potential vaccine expression platform. On the other hand, including the T2A element is beneficial for expressing foreign antigens as it allows co-expression of NSP3 and large peptides with little effect on the viral rescue efficiency, titre and replication, all which are important traits for live attenuated vaccine development. In the absence of the T2A element there may be a limit to the amount of additional foreign sequences that the RF NSP3 can accommodate. Our results offer a possibility of utilising a bovine RF RV as a backbone to facilitate the development of recombinant RV-based vaccines; here we used SARS-CoV-2 as an example, but this can be extrapolated to other gastrointestinal pathogens.

## Acknowledgements

We want to thank lab members, central support unit (CSU), bioimaging and technical staff for their support and assistance with this project. We thank the MRC-Protein Phosphorylation and Ubiquitylation Unit, University of Dundee and Dr Christine Tait-Burkard for providing SARS-CoV-2 spike antibodies and Mr Marius Diebold for various software support.

This study was supported by an ISSF3 Award from the Wellcome Trust. JS, PD and EG are supported by BBSRC Institute Strategic Programme Grant funding (BB/P013740/1) from the British Biotechnology and Biological Sciences Research Council). OD is supported by a Roslin Studentship Award. RB is supported by the University of Edinburgh scholarship. SC is supported by a Wellcome Trust Clinical Research Career Development Fellowship (211138/Z/18/Z). AB and EG are Sir Henry Dale Fellows supported by the Wellcome Trust (213437/Z/18/Z and 211222_Z_18_Z respectively). Funding for open access charge: Wellcome Trust.

We declare no conflicts of interest.

